# Wild type AAV, recombinant AAV, and Adenovirus super infection impact on AAV vector mobilization

**DOI:** 10.1101/2020.05.13.094201

**Authors:** Liujiang Song, R. Jude Samulski, Matthew L. Hirsch

**Affiliations:** Gene Therapy Center, University of North Carolina at Chapel Hill, NC, 27599, USA; Department of Ophthalmology, University of North Carolina, Chapel Hill, NC, 27599, USA; Department of Pharmacology, University of North Carolina at Chapel Hill, NC, 27599, USA

## Abstract

Recombinant Adeno-associated viral vector (rAAV) mobilization is a largely theoretical process in which intact AAV vectors spread or “mobilize” from transduced cells and infect additional cells within, or external, of the initial host. This process can be replication independent (vector alone), or replication-dependent (*de novo* rAAV production facilitated by super-infection of both wild-type AAV (wtAAV) and Ad helper virus). Herein, rAAV production and mobilization with and without wtAAV were analyzed following plasmid transfection or viral transduction utilizing well established *in vitro* conditions and analytical measurements. During *in vitro* production, wtAAV produced the highest titer with rAAV-luc (4.1 Kb), rAAV-IDUA (3.7 Kb), and rAAV-NanoDysferlin (4.9 Kb) generating 2.5-, 5.9-, or 10.7-fold lower amounts, respectively. Surprisingly, cotransfection of a wtAAV and a rAAV plasmid resulted in a uniform decrease in production of wtAAV in all instances with a concomitant increase of rAAV such that wtAAV:rAAV titers were at a ratio of 1:1 for all constructs investigated. These results were shown to be independent of the rAAV transgenic sequence, size, transgene, or promoter choice and point to novel aspects of wtAAV complementation that enhance current vector production systems yet to be de fined. In a mobilization assay, a sizeable amount of rAAV recovered from infected 293 cell lysate remained intact and competent for a secondary round of infection (termed non-replicative mobilization). In rAAV infected cells co-infected with Ad5 and wtAAV, rAAV particle production was increased > 50-fold compared to non-replicative conditions. In addition, replicative dependent rAAV vectors mobilized and resulted in >1,000 -fold transduction upon a subsequent 2^nd^ round infection, highlighting the reality of these theoretical safety concerns that can be manifested under various conditions. Overall, these studies document and signify the need for mobilization resistant vectors and the opportunity to derive better vector production systems.

## Introduction

Adeno-associated virus (AAV), a *dependovirus* of the family *parvoviridae*, was first identified as an Adenovirus (Ad) preparation contaminant in 1965 by Atchison *et al.* ^1^. The linear DNA AAV genome of approximately 4.7 kb consists of inverted terminal repeats (ITRs) flanking several open reading frames (ORFs), including *rep* and *cap*, which encode proteins involved in genome replication and capsid production respectively. Although AAV replication is not completely understood, the ITRs serve as the replication origins and work in concert with Rep proteins, the largest of which directly bind the ITR and induce a specific single strand nick as an initial step in the replication process^2,3^ Traditionally, AAV is considered a replication defective virus, that requires co-infection of a helper virus, several of which have been identified for completion of its natural life cycle^4^. In the absence of a helper virus, little expression of the *rep* ORFs occurs, and therefore, the AAV genome is minimally replicated and remains latent^5, 6^. However, AAV replication in the absence of a helper virus has been reported during cellular stress and/or in particular types of cells and/or phases of the cell cycle^7, 8^.

The wild type AAV2 (wtAAV2) genome was cloned into several plasmid constructs in the 1980s ^9–11^, and these constructs serve as the parental plasmids of most recombinant AAV (rAAV) vector constructs. In recombinant AAV (rAAV, also termed AAV vectors herein), the ITRs of serotype 2 (ITRs), are the only viral *cis* elements, flanking transgenic cassettes, as they are required for minimally, rAAV genome replication and capsid packaging^12^. Currently, AAV vectors are the most promising delivery method for *in vivo* human gene therapy with successes demonstrated in clinical trials for diverse diseases and a few drugs are FDA-approved and commercialized^2, 9, 10, 12–15^. Despite the popularity of AAV-based gene therapies, there remain unanswered questions regarding nearly all aspects of wtAAV and rAAV biology, in addition to the implications of the vector-induced genetic modifications in human patients^16^.

The rAAV vector itself is replication deficient, as replication requires the Rep proteins (absent from the vector), as well as a helper virus in most reported cases^10, 17–23^. However, wtAAV, which could supply the Rep and Cap proteins in *trans* for rAAV genome replication and capsid packaging, is prevalent in the human population^24^. Super-infection by other pathogenic viruses, such as herpes simplex virus (HSV) or Ad, which provide “helper functions” for wtAAV or rAAV, are also common in human patient populations. For example, the eye as an external organ offers unique advantages as a gene therapy target^25^, some of which also makes it more susceptible to viral infections. Previous reports demonstrated that herpetic keratitis, which is caused by HSV, is the most common corneal infection in the United States with 50,000 new and recurring cases diagnosed annually^26^. In addition, several Ad serotypes infect the conjunctiva and cornea and are responsible for 92% of all keratoconjunctivitis^27^.

rAAV mobilization is a largely theoretical process in which intact AAV vectors spread or “mobilize” from transduced cells and infect additional cells within, or external, of the initial host. This process can be replication independent, in which intracellular intact rAAV particles are released from the cell and infect another cell, or replication-dependent in which there is *de novo* rAAV production facilitated by super-infection of both wtAAV and a helper virus^28–32^. Despite nearly 40 years of rAAV investigations, including broad clinical applications, rAAV mobilization and its potential to induce disease and environmental safety concerns remains an overlooked and understudied problem with only a few publications investigating this phenomenon with no definitive quantification of the events ^29, 31^. In 1980, the rescue and mobilization of the wtAAV genome was first demonstrated in cell cultures^33^. In the case of rAAV, Tratschin et al reported a high frequency of integration and successful rescue from the chromosome by super infection of wtAAV and Ad, and in some cases, by infection with Ad alone in 293 and Hela cells in 1985^34^. Hewitt et al. demonstrated that rAAV vectors utilizing the ITR sequence from AAV5 reduce the risk of mobilization because of the lower frequency of wtAAV5 in human population combined with Rep2’s inability to nick the ITR5 sequence^29, 35^. Due to the lack of an appropriate animal model for studying the mobilization, so far, the only *in vivo* study to investigate this risk was carried out in non-human primates in 1996^31^. In that work, the authors demonstrated that rAAV replication and rescue occurred only after a direct administration of a very large dose of wtAAV into the lower respiratory tract prior to rAAV and Ad administration^31^. Whether this condition could ever happen in a natural setting is not clear, but illustrates the early efforts to address this hypothetical concern.

A more likely concern emanates from the risk of rAAV replication dependent mobilization released into the environment (or sheds) from treated patients following both systemic and local injections of AAV vectors ^36–41^. As shown in hemophilia clinical trials, AAV vector genomes were detected in the saliva, semen, blood serum, and urine of patients up to 12 weeks post intramuscular^42^ injections or portal vein infusion^43^. In clinical trials using Luxturna (which is locally administered directly to the subretinal space), rAAV shedding was observed in about 40% of the subjects through tears^44^. Although it remains to be tested if rAAV infectious particles are also shed from treated patients, these observations underscore the potential of shedding-associated rAAV transduction, with associated off-target transgenic DNA expression in unintended animal and human populations. Complicating this concern are studies, which demonstrate that capsid uncoating is a rate-limiting step in wtAAV and rAAV transduction^45–47^. Detection of intact rAAV particles by transmission electron microscopy up to 6 years after administration in the retinas of dogs and primates has been reported^48^. It remains possible that a large proportion of AAV vectors that fail to uncoat remain infectious^45, 49, 50^. Whether these vectors could be mobilized in a replication-independent manner and/or shed to mediate transduction in off-target cells, organs, and individuals remains unknown^29^.

Theoretically, during replication-dependent rAAV mobilization, AAV vector genomes compete with wtAAV genomes for the Rep and Cap proteins, which logically assumes that both wtAAV and rAAV production would decrease in the presence of a helper virus, if Rep or Cap proteins are rate-limiting for virion assembly. Herein, this theory is quantitively assessed under various conditions. The collective results demonstrate that following plasmid transfection: i) wtAAV production is more efficient than rAAVs, ii) rAAV titers vary significantly in a transgene-dependent manner, iii) all rAAV preparations herein are contaminated with wtAAV at a level of approximately 1%; however, following cotransfection of wtAAV and rAAV plasmids, wtAAV demonstrates a modest decrease in production titer while rAAV vectors are produced at increased levels following transfection; i) an effect also observed at the level of viral and vector genome replication. Following transduction, Ad co-infection dramatically increases rAAV transduction and replication-independent mobilization approximately 5-10 fold, ii) contamination of wtAAV in rAAV preparations is sufficient for replication dependent rAAV mobilization. iii) rAAV virion production was similar to wtAAV upon co-infection in the presence of Ad at a near 1:1 input ratio, and iv) replication-dependent mobilization resulted in ∼1,000-fold higher transduction compared to replication-independent mobilization. The data generated herein highlight the potential of rAAV vector production in treated patients, either receiving wtAAV in the vector preparation or upon subsequent wtAAV infection, superinfected by a helper virus and raise safety concerns for the treated individual and for the unintended animal and human populations in general.

## RESULTS

### Viral genome replication following plasmid transfection

To investigate an initial step during wtAAV and rAAV virion production, viral DNA replication was investigated following plasmid transfections in the absence or presence of the Ad helper plasmid pXX680^51^. Equal amounts of Hirt DNA isolated 48 hours (h) post-transfection were treated with *Dpn*I endonuclease to differentiate the input plasmids from newly replicated DNA, and then loaded for the agarose gel electrophoresis followed by Southern blot analysis. The wtAAV2 genome, the rAAV vector map, and the probes used for radioactive detection of the wtAAV or rAAV genomes are graphically depicted in Fig. 1A. A representative Southern blot revealed bands of the 4.7 kb monomer duplexes and the 9.4 kb dimer duplexes of wtAAV genomes, which indicated that the wtAAV2 genome was efficiently replicated following plasmid transfection with the presence of pXX680(Fig. 1B, lane ‘WT’). Under non-replicating conditions (without pXX680), wtAAV replication was not detected as expected (Fig. 1B, lane ‘Ctrl’). In the presence of a rAAV plasmid, the wtAAV genome was also replicated, however, to a lesser extent (Fig. 1B lane ‘WT’ vs. lane ‘WT+rAAV’). As show in Fig. 1C, both monomeric and dimeric rAAV-Luc replicative forms were observed in the rAAV only group (Fig. 1C, lane rAAV), and rAAV and wtAAV co-transfection group (Fig. 1C, lane WT+rAAV). Interestingly, the rAAV DNA was observed to replicate slightly more efficiently than in the absence of the wtAAV plasmid (Fig. 1C, lane ‘rAAV’ vs. ‘WT+rAAV’).

**Figure 1.**
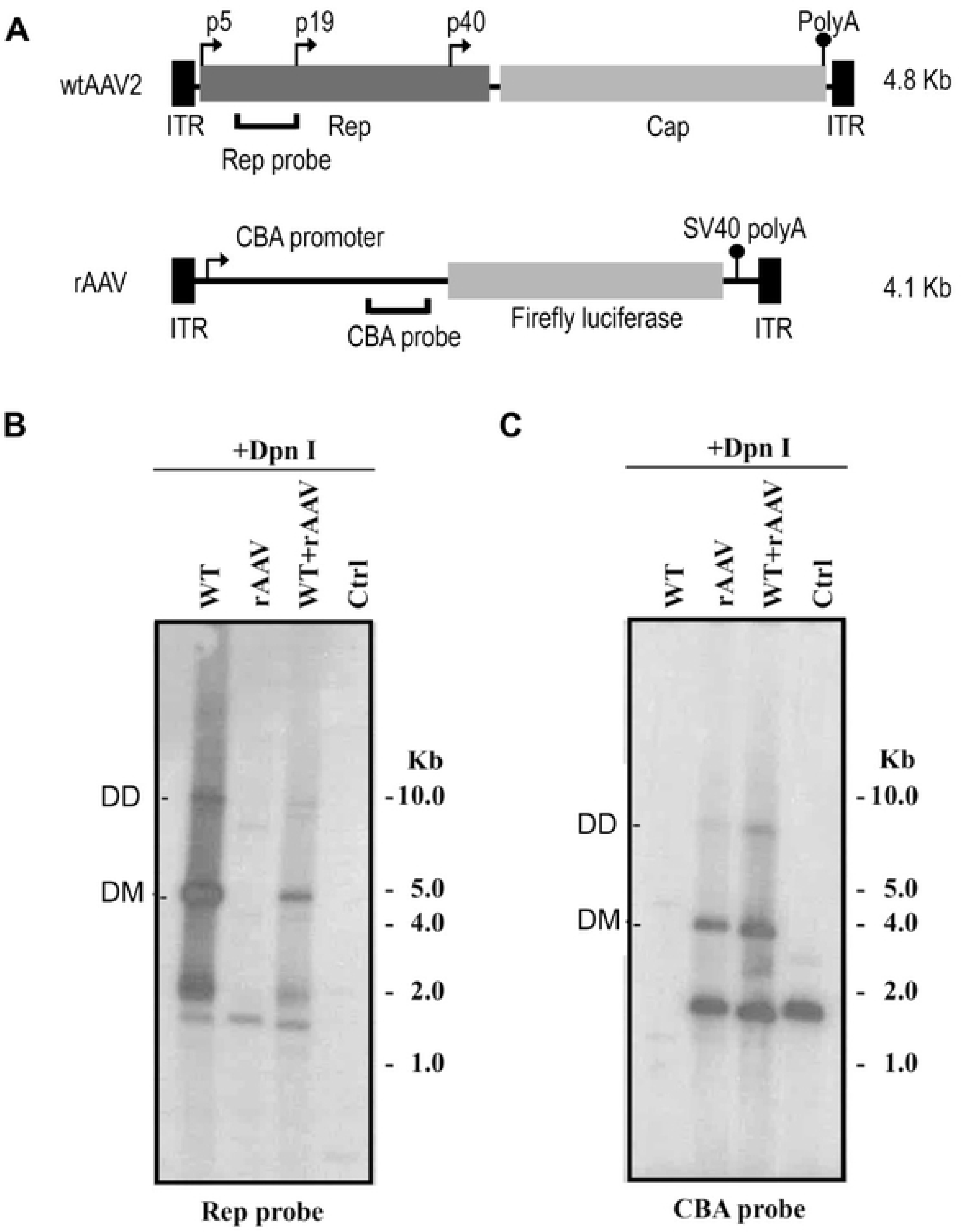
Replication assay of wtAAV and rAAV following cotransfection of wt and rAAV plasmids detected by Southern blot analysis. (A) Schematic maps of the wtAAV2 and rAAV-CBA-Luc including the probes used for detection. Southern blot analysis of wtAAV2 (B) or rAAV DNA replication (C) using the indicated probes. ITR, inverted terminal repeats; CBA, Cytomegalovirus enhancer/chicken β-actin promoter.

### AAV virion production following plasmid transfection

First, the wtAAV or rAAV particle yields were determined individually following a triple transfection protocol in 293 cells^51, 52^ using 15-cm-diameter plates. As described in the Methods, for rAAV production, a rAAV plasmid {pITR2-CBA-luc (4.1 Kb), or pITR2-EF1α-opt-IDUA (3.7 Kb)^53^, or pITR2-C2C27-Nano-dysferlin (4.9 Kb)^54^} was used along with pXR2 (*rep*2*cap*2)^51^ and the Ad helper plasmid pXX680^51^. To produce wtAAV, pSSV9^11^ was used along with pXX680 and an additional plasmid, pcDNA3.1 (to maintain consistent molar amount of total plasmids). Sixty-five h post-transfection, cells were lysed and virions were purified by CsCl gradient centrifugation. Alkaline gel electrophoresis followed by SYBR gold staining were used to visualize the packaged genome integrity and relative viral titer by loading the same recovered volume of the preparation (Fig. 2A), and quantitative PCR (qPCR) was utilized to determine the absolute titers (Fig. 2B). When packaging the wtAAV and rAAV separately, wtAAV2 preparations consistently demonstrated the highest titer at about 4.5e^11^ viral genome/plate (vg/plate) which was 3-10 fold greater than all rAAV2 preparations (Fig. 2B individual production groups). Among the three different rAAV2 preparations, significant differences in production titers were noted that varied approximately 3-4 fold, with AAV2-C2C27-Nano-dysferlin (4.9 kb) having the lowest titer of 4.2e^10^ vg/plate, AAV2-EF1α-opt-IDUA (3.7 kb) being intermediate at 7.6e^10^ vg/plate, while AAV2-CBA-luc produced the most rAAV2 particles at 1.8e^11^ vg/plate (Fig. 2B co-production groups).

**Figure 2.**
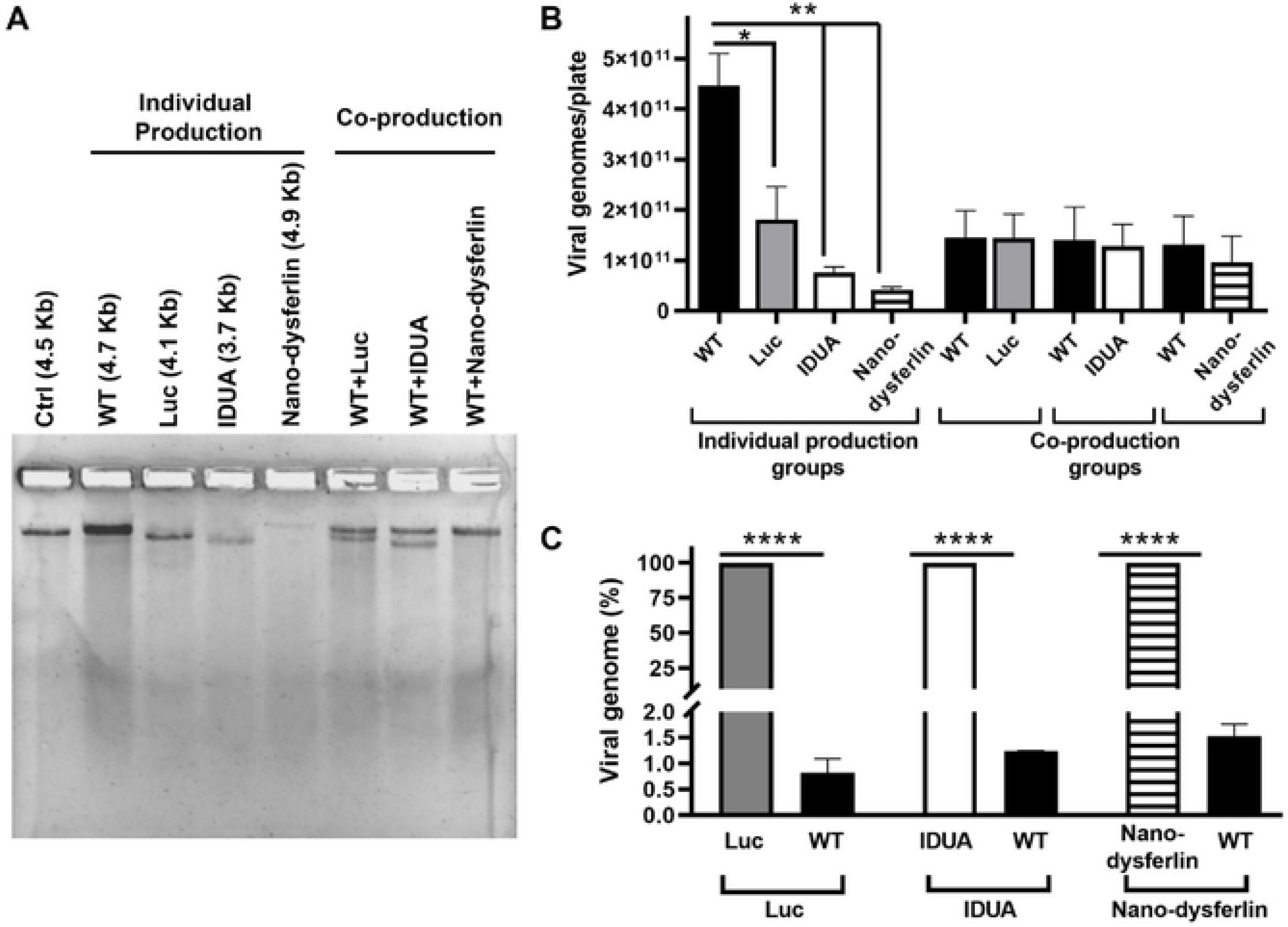
AAV virion production following cotransfection of wt and rAAV plasmids. (A) Viral genome integrity and relative viral titer visualized by alkaline agarose gel electrophoresis. A reference vector was loaded at 5e^10^ vg (Ctrl). (B) Virus titer determined by probe-based qPCR presented as the total vg/plate (mean± SD). *, p<0.05, **, p<0.01, Ordinary one-way ANOVA, Dunnett’s multiple comparisons test. (C) wtAAV contamination determined by qPCR in rAAV preparations. The results were normalized to each rAAV preparation and shown as percentage. ****, p<0.0001, Ordinary one-way ANOVA, Sidak’s multiple comparisons test. WT, wild type AAV2; Luc, AAV2-CBA-Luc; IDUA, AAV2-EF1α-IDUA; Nano-dysferlin, AAV2-C2C27-Nano-dysferlin.

In addition, wtAAV contamination in the three different rAAV preparations was investigated. The results of qPCR using a probe specific to *cap*2 sequence demonstrate that, when producing rAAV vectors with a triple plasmid transfection protocol in 293 cells, one of the most commonly used methods^51^, all rAAV preparations contained wtAAV particle contamination (or at least Benzonase-resistant *cap2* sequence), the amount of which ranged from 0.8-1.7% of the intended encapsidated rAAV genomes, depending on the transgenic sequence and/or size (Fig. 2C).

Next, rAAV and wtAAV virion production was assessed following plasmid cotransfections of wtAAV and rAAV plasmids. These experiments relied on the co-transfection of equal moles of pSSV9 and pITR2-transgenic, along with pXX680. In contrast to the production of wtAAV and rAAV seperately, wtAAV2 virion production during co-production setting was decreased approximately 3-fold with titers not significantly different than production of rAAV2 (Fig. 2B co-production groups). This result was shown to be independent of the transgenic sequence of the rAAV vectors used herein (Fig. 2B co-production groups).

### Significant replication-independent rAAV mobilization occurs

Replication-independent mobilization events following primary cell transduction was investigated by a secondary infection. Briefly, 293 cells were infected with rAAV2-CBA-Luc at doses of 100, 1,000 or 10,000 vg/cell. After 48 h post-infection, the medium was removed; cells were harvested and extensively washed with PBS. The cell pellets were then resuspended in 200 µl of PBS and subjected to freeze-thaw. Then the cell debris was removed by centrifugation, and the supernatant was used for qPCR detection of intact rAAV virons and for their ability to mediate subsequent transduction in fresh cells (Fig. 3). As described in the methods, for qPCR detection, the harvested supernatant was digested by Benzonase to remove nucleic acids that are not encapsidated, and then heated at 95 °C to denature the AAV capsid and to release the viral genomes. After 48 h post-infection, intact rAAV remained at detectable levels depending on the infectious dose (Fig. 3A). To check whether these intracellularly released rAAV virions were competent for replication-independent mobilization, the lysate supernatants from individual wells were added to fresh 293 cells for a 2^nd^ round of infection with or without co-infection by Adenovirus 5 (Ad5). Approximately 42 h post-addition, these 2nd round cells were lysed and subjected to luciferase activity detection to measure transduction. As shown in Figs. 3B and S1B, dose-dependent rAAV mobilization occurred resulting in luciferase activity. In the presence of Ad5, rAAV secondary transduction increased >10-fold compared to the no Ad5 groups (Fig. 3B). It should be noted that several controls were performed as outlined in the Methods to demonstrate that these results are not carryover of luciferase protein from the first round lysate and therefore, represent productive replication independent rAAV mobilization that is significantly enhanced by Ad5 following serial infection.

**Figure 3.**
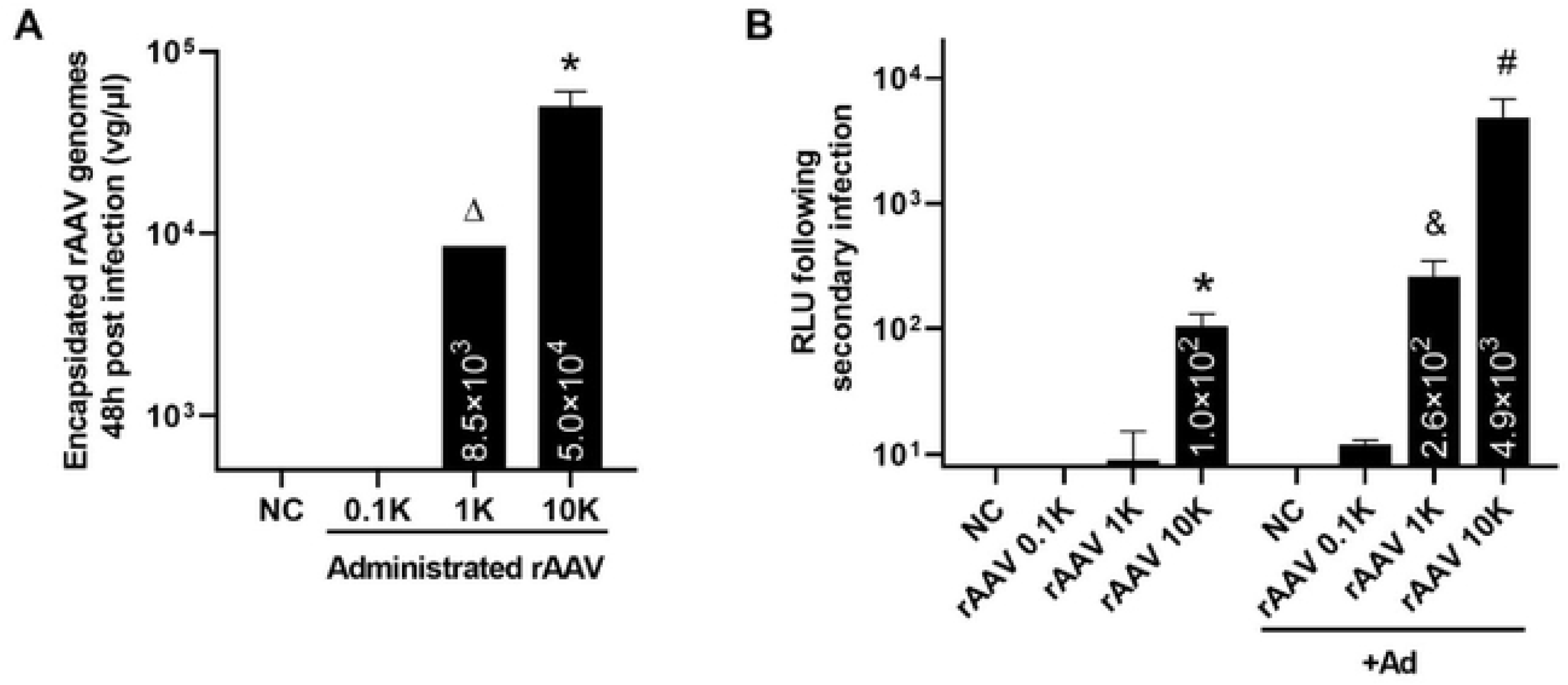
Replication independent-mobilization. (A) Quantitative analysis of the encapsidated rAAV using probe-based qPCR at 48 h post-infection. Cells were infected with rAAV at indicated doses. 48 h post-infection, cells were harvested, subjected to extensively wash and three rounds of freeze-thaw. Lysate debris was removed by centrifugation, and the supernatant was treated with Benzonase for 1h at 37 °C, heated at 95 °C for 15 min, diluted for at least 50 fold in H2o, and then used as a template for qPCR. The rAAV was detected using UPL probe/primers specific to the Luciferase coding sequence. Data are presented as mean ± SD (vg/µl). *, p<0.05. significant difference from NC. ∆, only 1 repeat was right above the lower detection limit, and the other repeats were detectable, but did not fall in the linear range of the standard curve.. (B) Luciferase activity from the 2^nd^ round of infection. Individual wells from the first round infection were subjected to extensively wash, then resuspended in the PBS, followed by freeze-thaw. The cell debris wasremoved by centrifugation, and the supernantant were added to fresh 293 cells with or without the presence of Ad for the 2^nd^ round infection. 42h post-infection, the cells were harvested for the luciferase activity measurement. Data are shown as mean±SD. *, p<0.05. significant difference from rAAV 1K, 0.1K and NC; ^#^, p<0.05, significant difference from the rAAV 10K (no Ad); ^&^, p<0.05, significant difference from the rAAV 1K (no Ad). Ordinary one-way ANOVA, N=3. NC, negative control without addition of AAV; rAAV, rAAV2-CBA-Luc; 0.1K, 1K and 10K represent rAAV was administrated at 100, 1,000 and 10,000 vg/cell, respectively.

### Significant replication-dependent rAAV mobilization occurs

Next, rAAV replication-dependent-mobilization in the presence and absence of wtAAV and a helper virus were investigated similarly by two rounds of infection. For the 1^st^ infection, 293 cells were infected with rAAV at increasing doses and/or wtAAV (fixed dose of 1,000 vg/cell). Ad5 (MOI=5) or PBS was added to the wells 4 h later. The cells were harvested 48 h post AAV addition and used for three different measurements: i) luciferase activity following the primary infection; ii) qPCR detection of encapsidated genomes; and iii) the 2^nd^ round transduction efficiency mediated by the rAAV released from primary infected cells at 42 h post-infection.

The luciferase activity of the first infection was measured at 48 h postinfection as described in Methods. As expected the rAAV transduction efficiency was dose dependent with dramatic enhancement in the presence of Ad5 (>10-fold increase) consistent with Fig. 3B and previous reports ^4, 55, 56^ (Fig. S1). However, when rAAV and wtAAV were co-administered to cells followed by Ad5 infection, luciferase activity in this setting decreased compared to the levels observed in the similar setting without wtAAV (Fig. S1).

To examine rAAV production in the presence of Ad5 and/or wtAAV in a transduction setting, qPCR was used to quantitatively assess the Benzonase-resistant (encapsidated) rAAV vector titer 48 h post-transduction. Interesting, in cells administered 1,000 or 10,000 rAAV vg/cell and Ad, without intentional wtAAV inoculation, the recovered rAAV titer increased approximately 2-5-fold in a dose-dependent manner (Fig. 4A, B, rAAV+Ad vs. rAAV). In cells given both rAAV + wtAAV (each at 1,000 vg/cell) and then super-infected by Ad5 (MOI=5), the encapsidated rAAV titer increased approximately 100-fold compared to transduction by rAAV alone (Fig. 4A, rAAV 1K+WT 1K+Ad vs. rAAV 1K) 48 h post-infection suggesting *de novo* rAAV production (Fig. 4A). This magnitude of this effect was observed to be dose dependent, in that when 10-fold more rAAV was used, only a 7-fold increase in rAAV titer was observed (Fig. 4B, rAAV 10K+WT 1K+Ad vs. rAAV 10K). When analyzing intact wtAAV production following transduction in cells super-infected with Ad5, a >6,000-fold increase in wtAAV virion titer was observed 48h post-infection (Fig. 4C, WT 1K+Ad vs. WT 1K). Interestingly, in cells co-infected with wtAAV and rAAV in the presence of Ad, wtAAV production was dramatically inhibited by rAAV in manner that directly correlated with the administered rAAV dose (Figs. 4C, S2). When the original input of rAAV was 10-fold higher than wtAAV, the total production of rAAV outcompeted the wtAAV (sFig. 2)

**Figure 4.**
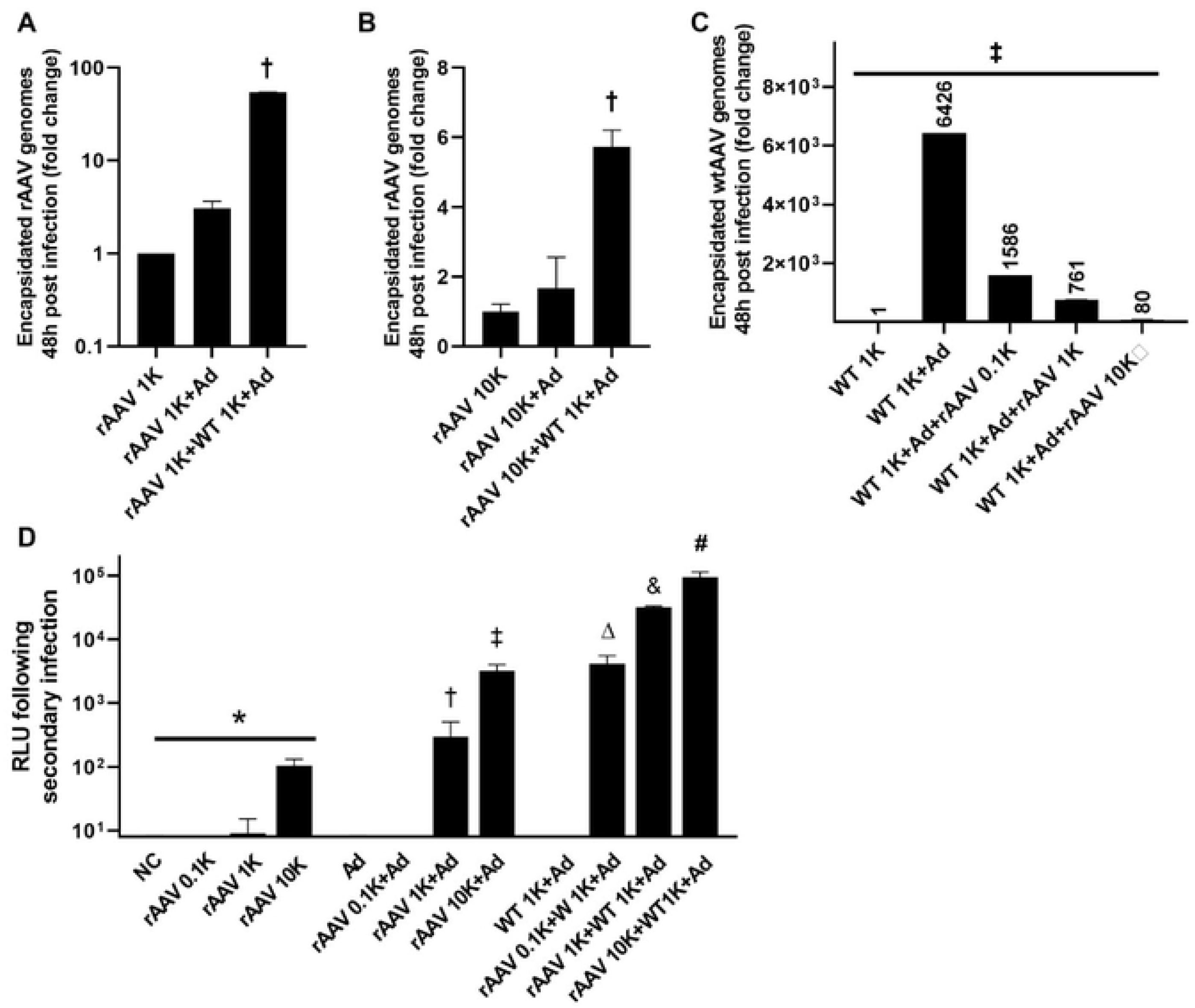
Quantitatively analysis of replication-dependent production and mobilization with the presence of wtAAV and or Ad. (A,B) Quantitative analysis of the rAAV vector genomes using probe-based qPCR. Cells were infected with rAAV at indicated doses, in the presence and absence of wtAAV (1,000 v/cell) and/or Ad (MOI=5). 48 h post rAAV infection, cells were harvested, extensively washed and digested with Benzonase. Encapsidated rAAV virions were detected by probe-based qPCR. Data are presented as mean±SD (vg/µl). †, p<0.001, significant difference from all; (C) Quantitative analysis of the wtAAV vector genomes using probe-based qPCR. Cells were infected with wtAAV (1,000 vg/cell), in the presence or absence of Ad (MOI=5) and/or rAAV2-CBA-Luc (at indicated doses). 48 h post-infection, cells were harvested, extensively washed and digested with Benzonase. Encapsidated wtAAV virions were detected by qPCR. Data were presented as mean±SD (vg/µl). ‡, p<0.0001; (D) Luciferase activity from the 2^nd^ round of infection. Cells from the first round infection were subjected to extensively wash, then resuspended inPBS, followed by freeze-thaw. The cell debris was removed by centrifugation, and the supernantant were added to fresh 293 cells for the 2^nd^ round infection. 42 h post, cells from the 2^nd^ round infection were harvested for the luciferase activity measurement. Data are presented as the mean±SD. RLU, relative luminesce unit. *, p<0.05, significant difference from the rAAV 1K, 0.1K and NC. #, significant different from rAAV 10K; &, p<0.05, significant difference from rAAV 1K; ∆, p<0.05, significant difference from rAAV 0.1K. NC, negative control without addition of AAV; WT, wtAAV2; rAAV, rAAV2-CBA-Luc; 0.1K, 1K and 10K represent AAV was administrated at 100, 1,000 and10,000 vg/cell, respectively.

Next, to determine if replication dependent mobilization occurs, the infected cell lysate was administered to fresh cells and rAAV transduction was assayed by luciferase activity 42h later. Significant enhancement of secondary infection was observed in primary cells administered rAAV and Ad with no intentional wtAAV administration (Fig. 4D, rAAV+Ad vs. rAAV). In primary cells co-administered rAAV and wtAAV in the presence of Ad, mobilized rAAV resulted in ∼1,000-fold enhancement of luciferase activity following secondary cell transduction depending on the original dose (Fig. 4D, rAAV+WT 1K+Ad vs. rAAV).

### wtAAV contamination in the rAAV preparations may facilitate the replication-dependent replication

wtAAV contamination of all rAAV preparations used herein was observed which is consistent with observations of multiple other groups^57, 58^ when using the triple transfection production protocol (Fig. 2C). We next examined whether these wtAAV contaminants could supply the required *Rep* and *Cap* for replication-dependent mobilization. The wtAAV and rAAV titer was examined after 48h post-addition of Ad5 (MOI=5). As shown in Fig. 5, the wtAAV production was increased more than 1,000-fold in the presence of Ad5 compared to the no Ad5 group. Interestingly, although the wtAAV sequence was not detectable in the rAAV-Luc alone groups, following super-infection of Ad5, detection of the wtAAV genome became evident in the 10,000 vg/cell dose, suggesting Ad-dependent replication and encapsidation of wtAAV contaminant particles in the rAAV preparation. This observation highly suggests that the contaminating wtAAV-like particles detected in rAAV preps were replication competent (Fig. 5).

**Figure 5.**
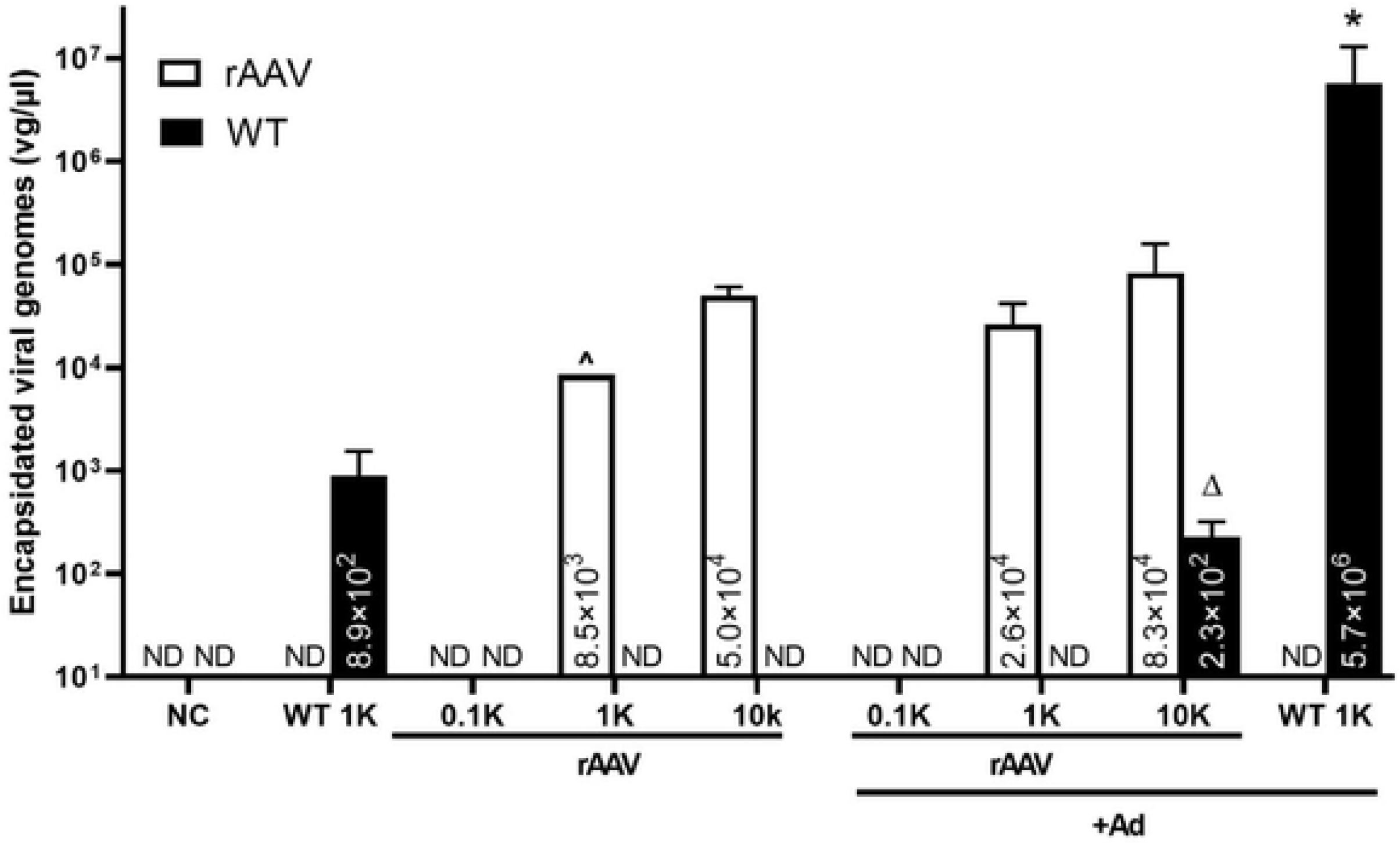
Quantitative analysis of replication-competent wtAAV. Cells were infected with wtAAV (1,000 vg/cell) or rAAV at indicated doses, with or without Ad (MOI=5). 48 h post-infection, cells were harvested, subjected to extensively wash and three rounds of freeze-thaw. The lysate debris was removed by centrifugation and the lysate supernatant was treated with Benzonase for 1h at 37 °C, heated at 95 °C for 15 min, diluted in molecular grade water and used for qPCR using UPL probes, the rAAV genome was detected with probe/primers specific to luciferase coding sequence, and the wtAAV genome was detected using probe/primers specific to *Cap* gene. Data are presented as the mean±SD (vg/µl). *, p<0.05, significant difference from WT 1K; Δ, statistics were impossible to be calculated due to the wtAAV genome in the rAAV alone groups being below the detection limit. ^, only 1 in 3 repeats were right above the detection limit.

In addition, the rAAV-Luc were slightly increased in both doses of 1,000 and 10,000 vg/cell, further suggesting that these replication-competent wtAAV might supply the *Rep* and *Cap* for the production of rAAV-Luc.

## Discussion

rAAV has become one of the most utilized delivery formats for human gene therapy with hundreds of clinical trials addressing numerous diverse diseases. In fact, currently, 3 rAAV drugs have been approved by the European and the U.S. Food and Drug Administration and estimations by the FDA have predicted that 10-20 new gene therapy drugs will be approved by 2025. Generally, positive therapeutic results have been obtained from clinical applications and overall, AAV gene therapy has garnered a strong safety profile. However, newly characterized concerns regarding the therapeutic use of AAV vectors remain including: 1) capsid and transgenic product immunogenicity, 2) demonstrations of chromosomal integration, 3) the potential for oncogenesis, and 4) potential complications related to off-target transduction in the treated subjects^59–64^ including non-patient bystanders resulting from rAAV shedding^37, 38, 40^. Surprisingly, rAAV mobilization, another largely theoretical concern associated with rAAV vectors, has been underappreciated in the AAV research community with only sparse reports over several decades^*10*, *29*, *31*, *33*^. wtAAV is reported to have better yields than rAAV during production in the laboratory^65^ yet, in a gene therapy setting, when a cell is transduced by rAAV and co- or super-infected by wtAAV and a helper virus, it is unclear whether wtAAV retains its advantage for replication and/or capsid packaging thereby potentially inhibiting de novo rAAV production in patients. In the current study, the reality of these concerns have been investigated in a quantitative manner in both transfection and transduction contexts in cell culture with several novel findings: i) replication-independent rAAV mobilization resulting in serial transduction is substantial (Figs. 3 and 6); ii) with the presence of both wtAAV and rAAV, there is no obvious bias between wtAAV and rAAV for production; however, a dose-dependent effect exists (Figs. 4C, S2); iii) wtAAV contamination of rAAV stocks facilitates rAAV *de novo* production and mobilization (Figs. 4A, B, and 5); and iv) replication-dependent mobilization results in >1,000 fold higher serial transduction compared to replication-independent mobilization (Figs. 4D, 6). The data generated herein expose the potential for AAV gene therapy treated patients to produce and disseminate rAAV, and thereby highlight the need for better vector production protocols that eliminate wtAAV contamination and safer, mobilization-resistant AAV vectors. Additionally, a yet to be defined aspect that increases rAAV production in the presence of a wtAAV plasmid is reported (Fig. 2)

**Figure 6.**
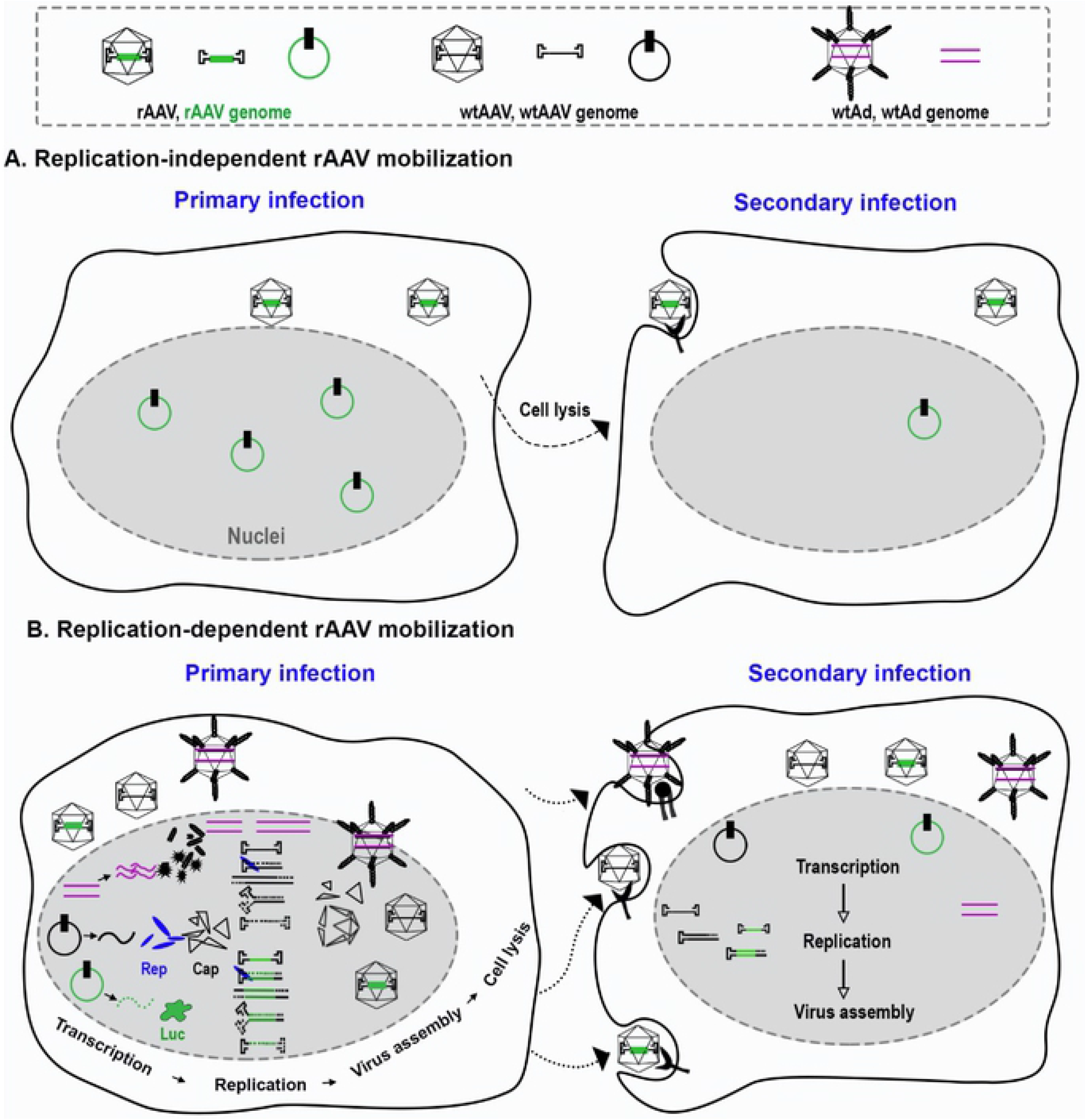
Model of replication-independent and -dependent rAAV mobilization. (A) Replication-independent mobilization. During the primary infection, some of the AAV vectors remain intact while others uncoat and persist in the nucleus as double-strand circular episomes that mediate transgene expression. Upon cell lysis, intact particles mobilize to other cells and mediate secondary cell transduction. (B) Replication-dependent mobilization. In the presence of wtAAV and a helper virus (Ad depicted), Ad provides helper function to the wtAAV to allow Rep and Cap gene expression. Rep proteins have the capacity to replicate both WT and transgenic AAV genomes and package them into AAV capsids resulting in de novo production of rAAV (and wtAAV) post-transduction. Upon cell lysis, the AAV virions mediate a 2^nd^ round of infection resulting in serial transduction by both previously uncoated, and newly produced, rAAV virion.

One obstacle for the characterization of rAAV mobilization is the lack of an ideal *in vivo* model to represent a natural infection situation in humans. In a lot of instances, Ad (or a different helper virus) and wtAAV present as natural co-infections^66^. It has long been the assumption that attributes of the wtAAV genome, such as the size and sequence, make it a more attractive substrate for Rep-mediated replication and capsid packaging compared to a rAAV transgenic genome^65^. This hypothetical preference for the wtAAV was observed previously^65^ and consistently herein, with wtAAV producing at a 3-10 folder higher titer compared to rAAV depending on the transgene (Fig. 2B). However, following co-transfection of wtAAV and rAAV plasmids, rAAV vector DNA tends to replicate more efficiently in the presence of wtAAV (Fig. 1C), and there is little to no bias observed toward wtAAV production (Fig. 2A, B). This may be attributed to the equal efficiency of wtAAV and rAAV genome encapsidation^67^, yet the improved efficiency of rAAV vector yield may be a result of when the *rep* and *cap* genes are supplied by the wtAAV genome, which is assumed to have the optimal *rep/cap* expression ratios during production in the laboratory setting^68^. Although still unknown, this result may be related to: 1) inherent ITR transcriptional activity that may fine-tune expression of known or currently undescribed AAV ORFs^69^ and or perhaps 2) the ability of the wtAAV helper plasmid to replicate, an activity shown to increase rAAV production^68^. Since the rAAV vector production via co-transfection in the laboratory mimics the production of wtAAV and rAAV with regard to replication substrates, these observations strongly suggest the likelihood of wtAAV and rAAV particle production at similar efficiencies post co-transduction (Fig. 2).

In the studies herein, it was also found that rAAV preparations produced by a common triple transfection protocol^70^ are contaminated with wtAAV particles (or at least Benzonase resistant wt capsid sequence) (Fig. 2C). This finding is consistent with several other reports demonstrating wtAAV contamination in rAAV preparations at a range of 0.01-10%^37, 57, 58, 71^. If these wtAAV particles proved to be replication competent virus, it decreases the stringency required for AAV vector mobilization, potentially by eliminating the requirement for subsequent (or prior) wtAAV transduction of a rAAV genome harboring cell (since it was administered to the patient as a rAAV preparation contaminant). The data show that upon Ad infection, the wtAAV-“like” particles significantly increased in the rAAV alone group, indicating that the contaminants are replication competent, consistent with a previous observation (Fig. 5)^58^. A number of approaches have been used to obtain replication-competent wtAAV-free stocks of rAAV^57, 58^, however, based on our observations, it is not clear if these approaches have gained widespread adoption for clinical rAAV production.

In the transduction analysis, rAAV transduction efficiency was substantially increased by the presence of Ad5 by an unknown mechanism, consistent with published studies^55^. Interestingly, the Ad enhancement appeared to be inhibited by the addition of wtAAV, since rAAV transduction in the presence of Ad and wtAAV was decreased when compared to rAAV+Ad groups (sFig. 2), this phenomenon is not understood and requires additional investigation^72, 73^.

Notably, rAAV particle production, following cell transduction, increased up to 100-fold in cells also infected with wtAAV and Ad (Fig.4A). These replication-dependent rAAV vectors, mobilized and resulted in a 1,000-fold increase in transduction efficiency in subsequently infected cells (Figs. 4D, 6). The extent of rAAV production directly correlated to the original input of materials, and the increased production of rAAV decreased the magnitude of the wtAAV production (Figs. 4C, S2). Although we did look at ratios of wtAAV to rAAV in the above experiment, it should be noted that wtAAV contamination in clinical rAAV preparations is likely several magnitudes less than the intended AAV vector (Fig. 2), thereby conferring a production advantage to rAAV in treated patient cells. This may lead to transgene specific toxicity as particular transgene products used in clinic could have detrimental effects in off-target cells and or healthy by-standers with the observations that tissue-specific promoter restriction and regulatory mRNA targets engineered into transgenic cassettes are incomplete.

Regarding the overall safety concern of rAAV mobilization, usually neutralizing antibodies to the AAV capsid inhibit super-infection of the same AAV serotype (dependent upon dose, route of administration, etc). The level of the antibody response to date from clinical trials has prevented re-administration and typically prevent all subsequent serotype administration due to cross antigen presentation. Although this immune response is extremely robust based on current vector doses, there is a window of vulnerability, namely: i) wtAAV infection post-AAV vector administration, yet prior to the onset of capsid neutralizing antibody production (an approximate 2 week window^74^), ii) super-infection of a wtAAV serotype that can evade the neutralizing antibodies induced by the therapeutic vector, iii) local injection of rAAV that results in minimal to no neutralizing antibody generation ^44, 53^, and iv) wtAAV as a contaminant of clinical rAAV preparations that is co-injected into the patient at the same time as the rAAV therapeutic and activated by helper virus or various stress conditions^37, 58, 71^. Concerns of inter-human spread of replication-independent and/or replication-dependent mobilized rAAV represents a formal safety concern for most mammals in general, and is theoretically dependent upon many factors including pre-existing capsid neutralizing antibodies, route of transmission, and helper virus and/or wtAAV infection in the mammalian bystander. However, the reality of this situation remains to be formally demonstrated.

In conclusion, the collective data emphasizes the pragmatic risk of rAAV mobilization in clinical settings from treated patients to the general public. Furthermore, evidence of a diminished capsid neutralizing antibody response over time in patients rekindles the risk of rAAV mobilization as well^75^. As the field progresses to more efficient AAV vectors for targeting, tropism, and ability to treat more broad diseases the data herein provided a proof-of-concept for safety concerns and highlight the need for mobilization resistant AAV vectors. One methodology is to alter the rAAV ITR sequence in a manner that allows efficient AAV vector production only in a laboratory setting. This notion was partially supported by a previous report in which vector genomes utilizing the rarer ITR5 sequence that are resistant to replication initiated by common Rep 2 or several other AAV serotypes were generated and tested, although wtAAV5 still exists in human populations^29^. Ultimately, rationally designed ITRs that are resistant to replication/mobilization by all naturally occurring Rep proteins yet allow high titer production by a novel laboratory restricted Rep-like protein would overcome rAAV mobilization concerns highlighted herein are required and such reagents are already under active investigation^35^.

## MATERIALS AND METHODS

### Cell culture

The HEK293 cells were maintained in 15-cm-diameter plates in Dulbecco’s modified Eagle medium (Gibco) supplemented with 10% heat-inactivated (30 min at 56°C) bovine calf serum (Gibco), 100 U of penicillin per ml, and 100 mg of streptomycin (Gibco) per ml at 37°C in a 5% CO2–air atmosphere.

### Plasmids

The plasmids used for transfections were as follows: i) The rAAV vector plasmids with the AAV2 inverted terminal repeats (ITRs)^9, 10^: pAAV2-CBA-Luc containing a firefly luciferase reporter cassette^70^, pAAV2-EF1α-IDUA harboring a clinical relevant α-L-Iduronidase expression construct^76^, and pAAV2-C2C27-Nano-dysferlin^54^ than encodes a truncated human dysferlin protein; ii) the AAV “helper” plasmid pXR2, harboring wtAAV2 *rep* and AAV2 *cap* sequence (without ITRs) ^52^; iii) The Ad helper plasmid, pXX680, encoding portions of the Ad genome that assist AAV production^51^; iv) the plasmid used for wtAAV production which encodes the AAV2 genome: pSSV9^11^; and v) pcDNA3.1 (originally obtained from Addgene, does not contain ITRs) was used to allow the same total molar mass to be the same for each transfection group.

### AAV DNA replication assay

The AAV DNA replication assay was performed using methods described previously with minor modifications^68, 77, 78^. Approximately 80% confluent 293 cells in 10-cm-diameter dishes were transfected with plasmids pXX680/pXR2/pAAV-CBA-Luc (to assess rAAV DNA replication), or pXX680/pSSV9/pcDNA3.1 (to assess wtAAV DNA replication), or pXX680/pSSV9/pAAV-CBA-Luc (to assess both rAAV and wtAAV replication), or pSSV9/pcDNA3.1 (serving as wtAAV DNA replication negative control), or pAAV-CBA-Luc/pXR2/pcDNA3.1 (serving as the rAAV DNA replication negative control). Fouty-eight h post-transfection, Hirt DNA was extracted. In short, the cell pellet from a 10-cm plate was re-suspended in 740 µl of Hirt buffer (0.01 M Tris-Hcl Ph 7.5 and 0.01 M EDTA) and lysed by adding 50 µl of 10% Sodium dodecyl sulfate (SDS). The cell lysate solution was then mixed with 330 µl of 5M NaCl, placed overnight at 4°C and centrifuged at 15,000 rpm at 4°C for 1 h. The supernatant was harvested and mixed with phenol/choloroform/isoamyl alcohol (25:24:1, Sigma) and centrifuged at maximum speed (∼16,000 rpm) for 5 min. The aqueous phase (top layer) was then mixed with an equal volume of chloroform (Sigma). Low-molecular weight DNA was precipitated by the addition of an equal volume of 100% isopropanol (Fisher), followed by centrifugation in a microcentrifuge tube at ∼16,000 rpm for 15 min at 4°C. The DNA pellet was dissolved in 50 µl of TE buffer (10 mM Tris–HCl, 1 mM EDTA pH 8.0) supplemented with 100 µg/ml of RNase A. The Hirt DNA yields were determined by the Nanodrop Spectrophotometer, and equivalent amounts of Hirt DNA, with or without prior digestion with *Dpn*I (New England Biolabs), were analyzed on 0.8% agarose gel followed by Southern blotting using a ^32^P-labeled DNA probe specific for AAV2 *rep* (for detection of wtAAV2), or CBA (for detection of rAAV2-CBA-Luc), prepared using a random primer labeling kit (Takara)^79^.

### AAV virus production and purification

AAV virions were prepared as previously described^77^, with minor modifications. Briefly, triple plasmid transfection of 293 adherent cells was performed with a polyethylenimine (PEI)-MAX (Polysciences, MW, 40,000) to DNA ratio of 3:1 (a mass-per-mass ratio). Equimolar of total plasmids was used for rAAV andwtAAV forvirus production.When packaging wtAAV and rAAV seperately, 0.8 mM (equal to 15 µg) of pXX680 plasmid, 1.6 mM of pXR2 plasmid (equal to 12 µg), and 1.6 mM of the rAAV plasmid were used per 15-cm-diameter dish of 80% confluent 293 cells for rAAV2 vector production. 0.8 mM of pXX680, 1.6 mM of pSSV9, and pcDNA3.1 were used for wtAAV2 production. Under coproduction condition, 0.8 mM of pXX680, 1.6 mM of pSSV9, and a rAAV plasmid were used for the transfection. pXX680 was co-transfected with pcDNA3.1 as a negative control. Cells and the growth medium were harvested around 65 h following transfection. The cell pellet was collected by centrifugation at 1500 rpm for 10 min. The virus in the growth medium was recovered by Polyethylene Glycol 8000 (PEG-8000, Fisher Scientific) precipitation. In short, 5x PEG/Nacl precipitation stock solution (containing 40% PEG-8000 plus 2.5 M Nacl in H2o) was added into the collected medium at a final concentration of 8% PEG followed by centrifugation at 5,000 rpm at 4°C for 30 min. The pellet was then resuspended with PBS and combined with the cell pellet and subjected to disruption by sonication. AAV virions were then purified by CsCl density gradient centrifugation^77^. The gradient fractions were then analyzed by alkaline gel electrophoresis and visualized by SYBR Gold Nucleic Acid Gel Stain (Thermo Fisher, Cat# S11494). The peak fraction(s) determined by the alkaline gels were combined, concentrated and desalted using Amicon Ultra-4 (100 K. MWCO, Millipore, Billerica, MA) with AAV storage solution (1× PBS with 5% sorbitol and 350 mM NaCl)^80^. Purified virus was bring to total volume of 1 ml in low retention tubes and stored at −80°C.

### Adenovirus

The human adenovirus type 5 (Ad5) stock was purchased from ATCC (VR-1516, Manassas, VA, USA) and employed as a helper virus for AAV replication^4, 51^. As a part of our initial characterization, the Ad5 infectious titer (Sfig. 3) were confirmed in 293 cells. Ad5 stocks were tittered by detection of the Ad hexon protein using Adeno-X™ Rapid Titer Kit (Cat. No. 632250, Takara, Clontech). The Ad5 CPE was dose-dependent, and at a MOI of 2 to 5 and cells showed obvious CPE after 48 h post-infection. Thus, for the experiments herein, a MOI of 5 was chosen unless otherwise indicated.

### AAV titer determination

wtAAV2 and rAAV2 particle numbers were determined by probe-based qPCR analysis following Benzonase treatment to eliminate non-encapsidated DNA^81^. Universal probe library (UPL) probe #3 (Roche) and following primers were used for analysis of wtAAV2: 5’-AGTACCAGCTCCCGTACGTC-3’ and 5’-CATACTGTGGCACCATGAAGAC-3’. UPL probe #29 and primers: forward, 5’-TGAGTACTTGAAATGTCCGTTC-3’ and 5’-GTATTCAGCCCATATCGTTTCAT-3’ were used for AAV2-CBA-Luc; UPL probe #15 and following primers were used for detection of AAV2-EF1α-IDUA: 5’-AAAGGGGGCCAGGTCTAGT-3’ and 5’-ATCTGCTGAGCGACCACCT-3’, UPL probe #46 and primers: forward 5’-CCGACACGCCTACCTGAG-3’ and reverse: 5’-CCGGCACTAAAATCGTCAG-3’ were used for AAV2-C2C27-Nano-dysferlin. Assay parameters with threshold cycle values and calculated quantities for each reaction were exported for further analysis. Plasmid standards were used to determine absolute titers.

### AAV mobilization detection

Replication-independent rAAV mobilization (defined as vectors failing to uncoat and shedding into non-targeted cells) and Replication-dependent mobilization of rAAV2 (defined as the rAAV vector genome being replicated and encapsidated in the presence of wtAAV2 and Ad5), was mimicked *in vitro* by successive infection of 293 cells. Briefly, 293 cells were seeded at 70% confluence in 24-well plates. The cultures were infected with rAAV2-CBA-Luc at either 100, 1,000 or 10,000 vg/cell, with or without the presence of wtAAV2 (1,000vg/cell) and Ad5. After 48 h post addition of Ad, the medium was removed, cells were harvested and washed with PBS. The cell pellets were then resuspended in 200 µl of PBS and subjected to three cycles of freeze-thaw lysis in a dry ice-ethanol bath. The cell debris was then removed by centrifugation, and the lysate supernatants were either used for qPCR detection of Benzonase resistant AAV genomes or for a second round infection. For the qPCR detection, equal volume of the lysate supernatant was digested with Benzonase for 1 h, heated at 95°C for 15 min, diluted in H_2_0 (due to the qPCR is sensitive to the inhibitors released from the crude lysate, the supernatant was diluted for at least 50 fold), and then directly used as a template for qPCR using UPL probes. The lower limit of detection (LOD, defined as the lowest titer detected within in the range of linear standard curve) of qPCR was determined according to the standard curve created by serial dilution of plasmids. To rule out pre-existing latent infection of wtAAV in the producer line, the 293 cells were routinely tested by qPCR and confirmed negative for wtAAV. Also, the plasmid preparations used for transfection, were all negative for wtAAV2 detection. For the second round infection, equal volume of the lysate supernatant was added to fresh 293 cells with or without addition of Ad (MOI=5). The transduction was evaluated approximately 42 h post-infection by reading the luciferase activity. In short, cells were harvested, extensively washed and directly lysed using 1X passive lysis buffer (Promega). Upon addition of substrate (Promega, E1483), light emission was measured using a Perkin Elmer plate reader, and the relative luminescence unit (RLU) were presented as the mean ±SD. Several controls were employed to exclude the possibility that the luciferase activity detected in the 2^nd^ round of infection was not carryover of luciferase protein from the first round cell lysate supernatant. First, as mentioned above, at the end of the 2^nd^ round infection, the medium, containing the lysate supernatant from the first round of cell lysates, was removed and the cells were extensively washed. In addition, when the lysate supernatant from luciferase plasmid transfected 293 cells was applied to fresh 293 cells, no luciferase activity was observed. Lastly, we further distinguished the luciferase activity by addition of Ad5 to all groups in the 2nd round of infection. Theoretically, the Ad5 would increase the rAAV transduction, but would be unlikely increase the activity of pre-existed luciferase protein carry over. The experiment was repeated on three separate occasions, and each experimental group was performed in triplicate.

### Statistics

Statistical analyses were conducted with Graphpad Prism software version 8 (San Diego, CA). Student’s t-test and Ordinary One-way ANOVA were used for data comparisons. Differences were considered significant when p < 0.05. Data are shown as mean ± SD.

## Acknowledgments

This study was supported by grants from the NIH RO1AI072176-06A1 (M.L.H.). Pfizer-NCBiotech Distinguished postdoctoral Fellowship (L.S.). The authors are grateful to Drs. Chengwen Li and Chuan-An Zhang for their excellent technique support. The authors thank Dr. Jacquelyn J. Bower, Prabhakar Bastola and Malik Moncalvo for their critical review of the manuscript.

---

**Supplementary Figure 1. Luciferase activity following primary infection.** Cells were infected with rAAV at indicated doses, with or without wtAAV (1,000 vg/cell) and/or Ad (MOI=5). 48 h post-infection, cells were harvested, subjected to extensively wash, then lysed and used for the luciferase activity measurement. Data are presented as the mean±SD. RLU, relative luminesce unit. *, p<0.05, significant difference from the Luc 10K. #, significant difference from Luc 1K; &, p<0.05, significant difference from Luc 0.1K. NC, negative control without addition of AAV; WT, wtAAV2; rAAV, rAAV2-CBA-Luc; 0.1K, 1K and 10K represent AAV was administrated at 100, 1,000 and10,000 vg/cell, respectively.

**Supplementary Figure 2. Relative fold change of encapsidated rAAV (normalized to encapsidated wtAAV) at 48h post-infection.** Cells were co-infected with wtAAV at a fixed dose (1,000 vg/cell) and rAAV at an escalating dose of 100, 1,000 and 10,000 vg/cell with the presence of Ad (MOI=5). 48 h post rAAV infection, cells were harvested, extensively washed and digested with Benzonase. Encapsidated AAV virions were detected using probe-based qPCR. The amount of encapsidated wtAAV at each group was set as 1, and the fold change of the encapsidated rAAV were normalized to the wtAAV at each dose. rAAV, rAAV2-CBA-Luc; 0.1K, 1K and 10K represent AAV was administrated at 100, 1,000 and 10,000 vg/cell, respectively.

**Supplementary Figure 3. Adenovirus tittering.** Infectious titer of Ad5 determined on 293 cells by detection of the Adhexon protein. Serial dilutions (10^−2^∼10^−8^) of the viral stock were used to infect HEK 293 cells. 48 h later, cells were fixed with methanol and stained with the antibody specific to the adenovirus hexon protein. Signal was detected after incubation with a secondary antibody conjugated with horseradish peroxidase (HRP) at room tempreture for 1h. Then the titer was determined by counting the number of brown cells (infected cells) in a given area. Each stained cell corresponds to a single infectious unit or event. The adenovirus infectious titer in this stock of Ad is 3e^10^ iu/ml.

